# Direct single cell-type gene expression analysis in Whole blood: Novel ratio-based gene expression biomarkers using 2 novel monocyte reference genes (*PSAP* and *CTSS*) for monitoring bacterial infection

**DOI:** 10.1101/2025.01.27.634977

**Authors:** Nelson LS Tang, Tsz-Ki Kwan, Dan Huang, Suk Ling Ma, K.S. Leung

## Abstract

**Background:** To determine single cell-type specific gene expression in peripheral blood (PB) requires either prior labour-intense cell sorting or expensive single-cell RNA sequencing. We developed and validated a novel ratio-based biomarker (RBB) called Direct Leukocyte Subpopulation-Transcript Abundance (DIRECT LS-TA) assay that allows quantification of monocyte-specific gene expression directly from peripheral blood samples without cell sorting.

**Methods:** The DIRECT LS-TA method leverages known cell-type proportions and differential gene expression profiles among leukocyte subpopulations (e.g. monocytes, lymphocytes and granulocytes) to identify monocyte-informative genes. We shortlisted genes that had 2.5-fold higher expression in isolated monocytes compared to PB samples, indicating >50% transcript of genes in PB are contributed by monocytes alone. Public gene expression datasets were used to generate a list of monocyte informative genes with which DIRECT LS-TA assay is applicable in PB samples. *PSAP* and *CTSS* were identified as the monocyte informative reference genes, based on low biological variation (CV <12%) and high monocyte specificity. They were used as the denominator and together with another monocyte informative target gene as the numerator, the value of this new DIRECT LS-TA can be determined which is a kind of ratio-based biomarker (RBB). The clinical utility was the differentiation of patients with bacterial infection from control subjects in a discovery dataset and 4 other replication datasets. Methods to convert DIRECT LS-TA results to multiple of control median (MoM) provided approximations to delta-delta CT values of relative quantification commonly used in qPCR or digital PCR assays.

**Results:** Over 50 monocyte-informative genes were identified, including key immune response genes like *IFI44L*, *IL1B*, *VNN1* and *NFKBIZ*. DIRECT LS-TA results showed excellent correlation with the gold standard results, gene expression in isolated monocyte expression (R^2^=0.55-0.97). Then, DIRECT LS-TA of these 50 target genes were evaluated to identify the best RBB to detect the host response to bacterial infection. Monocyte DIRECT LS-TA *VNN1* RBB showed consistent upregulation across five independent datasets (median fold change 2.7-fold, p<10^-8^) with strong diagnostic performance (AUC=0.84-0.99). The expected corresponding delta-delta CT value is more than 1.4 cycle which can be reliably measured by qPCR. Additional monocyte-informative genes including *NLRC4*, *CYP1B1*, *PFKFB3*, *LILRA5*, *NFKBIA*, and *NFKBIZ* also demonstrated significant diagnostic ability (AUC>0.8).

**Conclusions:** The DIRECT LS-TA method (as a simple RBB) provides reliable quantification of monocyte-specific gene expression directly from whole blood samples without cell separation. The robust performance in bacterial infection diagnosis demonstrates its potential clinical utility for rapid infection differentiation and informed antibiotic stewardship. DIRECT LS-TA will emerge as a new kind of in vitro diagnostics (IVD) which can convey single cell-type gene expression information from PB samples. The new kind of IVD and uniqueness of the information, together with the ease of implementation will make it very useful in clinics

## 1. Introduction

Expression levels or transcript abundance (TA) of genes in peripheral blood cells serve as important biomarkers. Current clinical biomarker applications of gene TA quantification in peripheral blood (PB) samples are performed on cell mixture samples such that the TA results represent summation of TA of all cell-types of leukocytes. Therefore, no information of TA of a specific leukocyte cell-type (leukocyte subpopulation, LS, such as B lymphocytes or monocytes) can be obtained from the bulk RNA quantification in PB.

On the other hand, TA of genes in specific LS are the preferred biomarkers. However, in order to obtain TA in a specific LS, the current state of the art requires prior cell sorting to isolate the specific LS from PB before quantification of TA by quantitative PCR (qPCR) or digital PCR (dPCR). Latest technology of single-cell RNA sequencing can also obtain TA of a specific LS even at the cellular level or TA of every single cell (Stephenson et al. 2021; Wang et al. 2021b). However, both methods are either not applicable or affordable in common clinical use. Cell sorting/isolation is too laborious and tedious to run in a routine hospital laboratory. Single-cell RNA sequencing is too expensive to use for every patient admitted for investigation of febrile illness, for example. These procedures are basically not practical in the setting of a clinical service laboratory. A straight-forward laboratory protocol to obtain TA for genes of interest of specific LS in PB is needed. We described a prototype of direct method to obtain single cell-type specific TA using B lymphocytes as the target(Tang 2017; Huang et al. 2021). Here, the same logistic is applied to target monocyte with the objectives of gathering TA in monocytes directly from PB without prior monocyte separation.

PB is composed of various key LS, including lymphocytes T cells, B cells, monocytes and natural killer cells (in PBMC) and granulocytes in addition in whole blood. They are present in different but within a given range of cell proportions. For example, B lymphocytes typically comprise about 5% of leukocytes in peripheral whole blood (WB). Monocytes accounts for 10% to 30% of leukocytes in PBMC. The direct single cell-type specific transcript abundance assay (DIRECT LS-TA) utilises the given cell proportion of the specified cell-type to shortlist a list of genes whose transcripts in PB are produced predominantly by that specified single cell-type. We have succeeded in using B cells TA response as early biomarker of seroconversion after vaccination in which a figure of 5% was used as the given B cell proportion in PB (Huang et al. 2021). With the proof of principle method development for cell proportion figure as low as 5%, this DIRECT LS-TA approach should be even more powerful with LS cell-types present in higher cell proportion. Monocytes are present at a higher cell proportion in PBMC and a figure of 20% was used here. In the method section, a more detailed description of how to shortlist monocyte informative genes are provided.

Other methods have developed for microarrays or bulk RNA-sequencing data of PB when TA of all genes are obtained. For example, deconvolution methods (Newman et al. 2015; Monaco et al. 2019; Avila Cobos et al. 2020) have been developed to determine the cell count proportion of each LS cell-type in a PB cell-mixture sample but most of them cannot obtain gene TA in an individual sample as TA of a gene is commonly assumed to fixed group-wise values for patients and controls groups. These deconvolution approaches fail to capture actual gene TA variations for an individual and require TA data of the whole genome generated from microarrays or bulk RNA-sequencing. Recently, there is a keen effort to obtain single-cell-type specific gene expression for individual samples in bulk RNA sequencing data of cell-mixture samples using AI assisted and Bayesian statistical methods (Wang et al. 2021a; Khatri et al. 2024; Chen et al. 2022; Newman et al. 2019; Chu et al. 2022; Fan et al. 2021). Again, these methods require input of TA data of the whole genome. In this study, we aimed to develop monocyte DIRECT LS-TA and obtain TA of monocytes in patients with bacterial infections.

## 2. Materials and Methods

### 2.1 Datasets Used in the Analysis of Gene Expression of Peripheral Blood and Monocytes

In order to identify monocyte informative genes that are suitable for the Direct LS-TA assay, the following gene expression datasets obtained from peripheral blood samples were used (Table 1). These datasets were available from the Gene Expression Omnibus (GEO), maintained by the US National Institutes of Health. Details were available under their accession numbers. The types of peripheral blood samples obtained included whole blood (WB) and peripheral blood mononuclear cells (PBMCs). Specific cell types that have been further isolated and purified, such as isolated and purified monocytes, were also included in some datasets.

**Table 1.** List of PBMC or WB gene expression datasets used to identify monocyte informative genes.

| Dataset accession number | Usage | Type of peripheral blood sample (WB or PBMC) | References |
| --- | --- | --- | --- |
| GSE138746 | 1. To calculate the 90<sup>th</sup> percentile value of the fold of expression in monocytes vs. PBMCs, and find out genes that meet the criteria for monocyte informative gene;<br>2. To examine and determine the correlation between the biomarker of gene expression obtained by using Direct monocyte LS-TA and the gene expression of isolated and purified monocytes (the gold standard) | PBMC and isolated and purified monocytes | (Tao et al. 2021) |
| GSE114407 | Ditto | PBMC and isolated and purified monocytes | (Kuan et al. 2019) |
| GSE60424 | Ditto | Whole blood and isolated and purified monocytes | (Linsley et al. 2014) |
| GSE107011 | To compare the gene expression of granulocytes and monocytes | PBMC and isolated and purified monocytes and granulocytes | (Monaco et al. 2019) |

### 2.2 Datasets Used in the Analysis of Gene Expression Markers of Monocytes for differentiation of bacterial infection

In order to identify monocyte marker that enables differentiation of bacterial infection, the following gene expression datasets obtained from peripheral blood samples were used (Table 2). GSE154918 is a RNA-sequencing dataset of PB of infection patients. It is used as the discovery dataset as RNA-sequencing results have a better coverage of the transcriptome and are not restrictive by the probes availability as in microarrays. After identifying potential RBB (monocyte DIRECT LS-TA) that were activated in monocyte after bacterial infection, these RBBs were evaluated in 4 other replication datasets. They were also PB gene expression datasets but were analyzed on microarray platforms (Illumina and Affymetrix). Different gene expression platforms were included here to show that the new monocyte DIRECT LS-TA was a genuine phenomenon of monocyte and was not analysis platform dependent. In all datasets, only results from patients with uncomplicated bacterial infection were used, while results from patients with sepsis (if any) were filtered off. Sepsis is a complication or dysfunction of the immune system towards heterogeneous insults and not only against bacterial infection.

**Table 2.** List of WB gene expression datasets used to identify monocyte DIRECT LS-TA biomarkers for differentiation of bacterial infection.

| Dataset accession number | Grouping of samples and number of samples | Type of blood sample<br>(WB or PBMC) | References |
| --- | --- | --- | --- |
|  | Discovery Dataset |  |  |
| GSE154918<br>(bulk RNA-seq) | Bacterial infection group: 11<br>Control group: 40<br>(samples from sepsis or follow-up patients were not included) | WB | (Herwanto et al. 2021) |
|  | Replication Datasets |  |  |
| GSE40012<br>(Illumina microarray) | Bacterial infection group: 30<br>(samples from patients in bacterial infection group on Day 1 and Day 2)<br>Control group: 36 (samples on Day 1 and Day 5) | WB | (Parnell et al. 2012) |
| GSE42026<br>(Illumina microarray) | Bacterial infection group: 18<br>Control group: 33 | WB | (Herberg et al. 2013) |
| GSE60244<br>(Illumina microarray) | Bacterial infection group: 22<br>Control group: 40 | WB | (Suarez et al. 2015) |
| GSE63990<br>(Affymetrix microarray) | Bacterial infection group: 71<br>Control group: 89 | WB | (Tsalik et al. 2016) |

### 2.3 To define Monocyte Informative Genes Whose Expression Level Can Be Reliably determined from Cell Mixture Samples e.g. PBMC, WB

In our previous publication, we defined cell-type informative genes as genes which are predominantly expressed by only a single cell-type (e.g. monocyte) to the extent that ≥50% of gene transcripts of these informative genes in a cell-mixture sample (for example, PBMCs) were contributed by that single cell-type (Huang et al. 2021). Typically, the cell count percentage of monocytes in PBMCs was 10%-30% (Meskini et al. 2024; Nielsen et al. 2020). The proportional cell count of monocytes in PBMCs was pre-defined to be 20% in this case. By using the definition we described previously(Huang et al. 2021), when the proportional cell count of monocyte was 20%, the expression of an informative gene in the monocyte sample needed to be 2.5 times higher than that in the cell-mixture sample as illustrated in Figure 1. The monocyte informative genes in the cell-mixture blood sample were identified by using these conditions. Expression data from the isolated monocyte sample and cell-mixture sample (PBMCs or WB) in datasets from GEO (Table 1) were used to determine which genes were monocyte informative genes.

**Figure 1.**
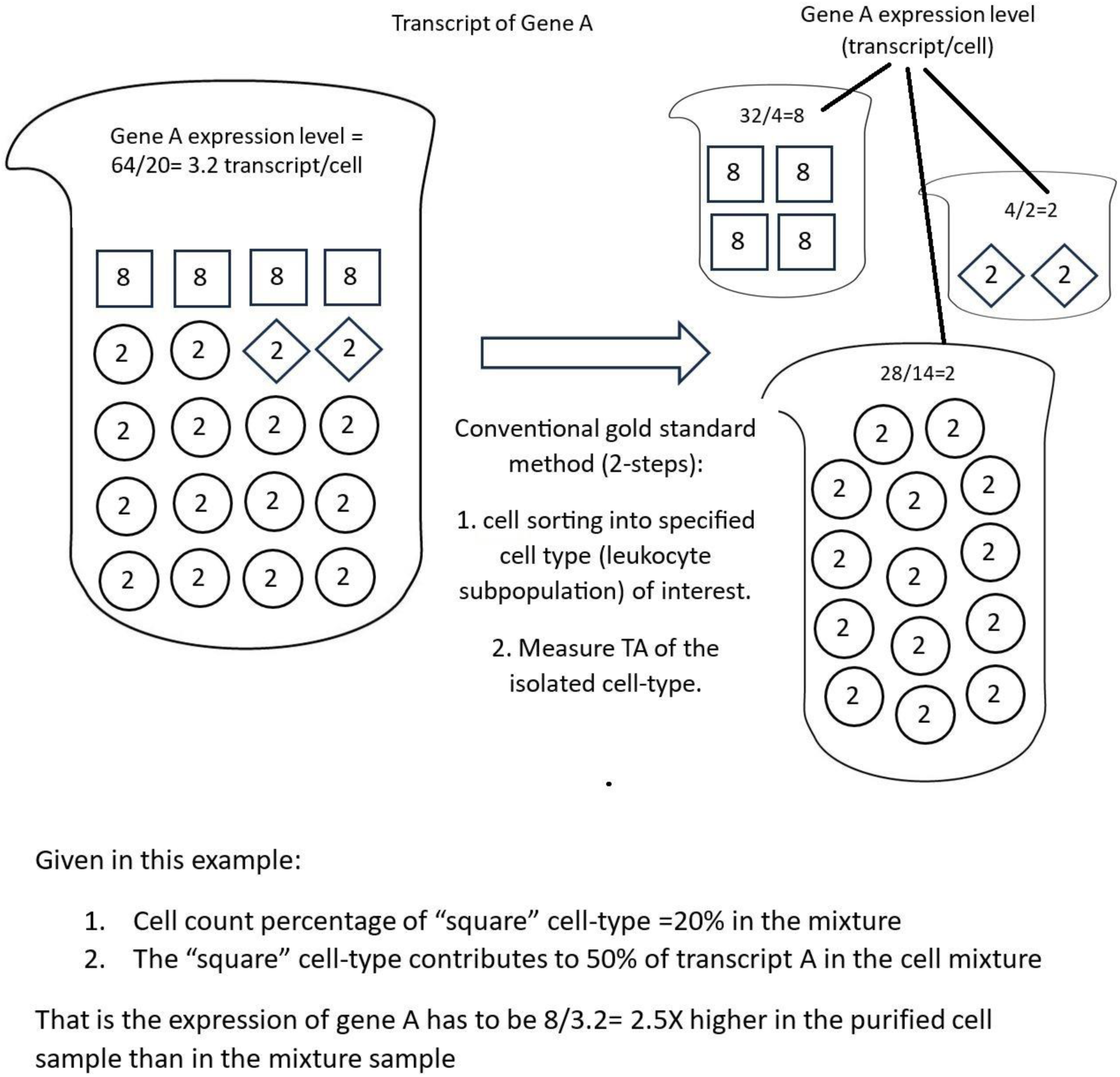
A schematic diagram of screening for cell subpopulation informative genes. The figure shows 4 samples, on left side is the original cell-mixture samples (e.g. PB) and 3 constitutional cell-types samples obtained from the original cell by cell separation / sorting, on the right side. Monocyte, the cell-type of interest, is shown as cells in square symbols. A pre-defined cell count percentage of monocyte is set to 20% as in PBMC. If any gene, such as gene A, has an average cellular expression level in an isolated monocyte sample that is above 2.5x (folds) higher than its average cellular expression level in a cell-mixture sample (for the square cell subpopulation, 8 (for isolated square cells) / 3.2 (in cell-mixture sample) = 2.5 folds), this cell subpopulation (i.e., the square cell subpopulation) contributes 50% of gene A transcripts in the cell-mixture sample. The gene expression level is represented by a ratio of number of transcripts to the number of cells. For example, the cell-mixture sample has a total of 64 transcripts of Gene A in 20 cells, the gene expression value = 64 / 20 = 3.2 transcript/cell. Such gene expression values are similar to the concept of relative expression quantification using housekeeping gene to normalize target gene expression. Genes with expression level 2.5x (folds) higher in the isolated square cell subpopulation than in the cell-mixture sample are potential monocyte informative genes. In this example, to demonstrate the principle, it is assumed that the gene expression levels of other cells are known. In fact, we only need to know the expression levels of the specified cell sample after isolation and the corresponding cell mixture sample to identify the single cell-type informative gene.

### 2.4. Development of DIRECT LS-TA value from gene expression datasets

To develop the RBB (DIRECT LS-TA), the denominator genes are required, they can be selected among the 50 shortlisted monocyte informative genes. The denominator gene (monocyte informative reference genes) are those monocyte informative genes with the least biological variation. Therefore, the percentage coefficient of variation (CV%) was calculated for each monocyte informative genes to find those with lowest CV%. Conventional housekeeping genes cannot be used here as they are expressed across all cell-types in PB but not specific to monocyte. We also evaluated if the typical cell marker gene for monocyte in used flow cytometry (*CD14*) could serve as the denominator (monocyte informative reference genes).

### 2.5 Devising the new monocyte cell-type specific RBB, monocyte DIRECT LS-TA from bulk transcriptome results of peripheral blood samples

Suppose both the numerator and denominator genes of the new RBB (DIRECT LS-TA) are only expressed by monocytes, the same ratio will be obtained no matter whether they were quantified in PB (cell-mixture sample) or the corresponding isolated monocyte sample. As genes with such degree of cell-type restrictive expression are few, we relax the degree of cell-type predominant expression to any genes with more than 50% of transcripts in PB come from monocytes. With such degree of preferential expression, the new RBB of PB formed by the ratio of a monocyte informative target gene over a monocyte informative reference gene (DIRECT LS-TA) may correlate with expression of that target gene in isolated monocytes. Therefore, DIRECT LS-TA values of all shortlisted monocyte informative genes were derived from the bulk gene expression results in cell-mixture samples (e.g. WB or PBMC) in datasets listed in Table 1 and their correlation with gold standard results (gene expression in purified monocytes) were analyzed. R^2^ of above 0.5 (r>0.7) is used as a cutoff of acceptable correlation and these target genes are then selected for further evaluation of potential RBB.

Monocyte DIRECT LS-TA = monocyte informative gene _(WB)_ / monocyte reference gene _(WB)._

That is the new RBB is a ratio of TA of 2 genes in the cell-mixture sample (e.g. WB or PBMC).

In the gene expression dataset, the gene expression values are log transformed. Therefore, this RBB can also take its log form as follows:

Log (monocyte DIRECT LS-TA) = Log (monocyte informative gene _(WB)_) - Log (monocyte reference gene _(WB)_)

Conceptually, this is similar to the results of delta CT (ΔCT) in qPCR relative quantification experiments. The difference of threshold cycles (CT) of target gene and normalisation reference gene (typically one or more housekeeping genes) is called delta CT (ΔCT) in qPCR relative quantification. In order to understand the degree of activation after stimulation or having a disease, such results (delta CT, ΔCT) can be compared to the delta CT of a control (or calibrator) individual, this new result is called delta-delta CT (ΔΔCT).

To reflect the extent of gene activation compared to the controls, a result that is conceptually similar to delta-delta CT (ΔΔCT) can be obtained for DIRECT LS-TA. The median DIRECT LS-TA of the control group is used as the calibrator individual, and it is subtracted from the value of DIRECT LS-TA of all subjects. In other word, the median DIRECT LS-TA value was set to zero and all other DIRECT LS-TA results were re-calibrated. Statistically, it represents a conversion of raw DIRECT LS-TA to the correspondent multiples of median (MoM). The median is defined from the control group of each dataset. The MoM results will be comparable to ΔΔCT results when DIRECT LS-TA is used in prospective patients with qPCR or dPCR assays.

Since experiments performed with different detection assays would yield results in different units, a method was needed to normalize the results across various datasets obtained from different detection methods. MoM was often used for assays that had not been standardized through large-scale assays, for example prenatal biochemical screening (Driscoll et al. 2009), and may be used in cytokine assays for determining the risk of adverse outcomes after SARS-CoV infection (Tang et al. 2005). The advantage of using MoM was that it may remove the limitation between datasets due to different measurement units among various laboratories, so that comparison may be performed on the results obtained by different laboratories or assays.

### 2.6 Data Analysis and Statistics methods

In the discovery dataset, GSE154918, monocyte DIRECT LS-TA results of 50 monocyte informative genes (see Table 3) were calculated. And the difference of monocyte DIRECT LS-TA result of each target gene was compared between the control group (n=40) and uncomplicated bacterial infection group (n=11). A non-parametric statistic (Wilcoxon–Mann–Whitney test) test was used to derive the p value. As there are 50 target genes and 2 reference genes generating 100 monocyte DIRECT LS-TA results for each sample, a multiple testing correction by Bonferroni method was used and the type I error is set to 1 x 10^-5^ (equivalent to corrected p<0.001).

**Table 3:** List of Monocyte cell-type specific Informative Target Genes and 2 Reference Genes (*CTSS* and *PSAP*) **\* *CTSS* and *PSAP* are used as denominator genes in the RBB, so they do not have correlation results to compare with gold standard method.**

| Reference and Target gene | The 90<sup>th</sup> percentile value of the fold of expression in monocytes vs. peripheral blood or PBMCs | Coefficient of variation (CV%) | Correlation between the Monocyte DIRECT LS-TA biomarker obtained directly from peripheral blood samples and the gold standard (expression level of target genes in isolated and purified monocytes) (coefficient of determination, $r^2$) | Database source showing the best correlation |
| --- | --- | --- | --- | --- |
| *CTSS* | 2.8x | 9% | NA* | GSE138746 |
| *PSAP* | 3.2x | 11% | NA* | GSE138746 |
| *ASGR2* | 3.6x | 41% | 0.79 | GSE138746 |
| *ATF3* | 3.9x | 164% | 0.64 | GSE138746 |
| *CALHM6* | 2.7x | 78% | 0.78 | GSE138746 |
| *CD163* | 3.5x | 65% | 0.85 | GSE138746 |
| *CD36* | 3.5x | 22% | 0.64 | GSE138746 |
| *CDKN1A* | 2.8x | 101% | 0.81 | GSE138746 |
| *CES1* | 3.7x | 51% | 0.85 | GSE138746 |
| *CLEC12A* | 3.1x | 33% | 0.73 | GSE138746 |
| *CRISPLD2* | 3.8x | 31% | 0.72 | GSE138746 |
| *CXCL10* | 3.6x | 212% | 0.84 | GSE60424 |
| *CYP1B1* | 3.9x | 45% | 0.80 | GSE138746 |
| *CYP27A1* | 4.1x | 30% | 0.65 | GSE138746 |
| *EREG* | 6.6x | 172% | 0.66 | GSE138746 |
| *FOS* | 4.3x | 83% | 0.82 | GSE138746 |
| *GADD45B* | 2.6x | 69% | 0.70 | GSE138746 |
| *IER2* | 3.0x | 80% | 0.74 | GSE138746 |
| *IFI30* | 3.3x | 20% | 0.60 | GSE60424 |
| *IFI44L* | 3.2x | 141% | 0.95 | GSE60424 |
| *IFITM3* | 2.7x | 100% | 0.87 | GSE138746 |
| *IL1B* | 3.8x | 183% | 0.80 | GSE138746 |
| *KLF10* | 2.6x | 52% | 0.83 | GSE138746 |
| *LILRA5* | 3.4x | 23% | 0.55 | GSE114407 |
| *LYZ* | 3.4x | 23% | 0.69 | GSE138746 |
| *MAFB* | 4.1x | 65% | 0.73 | GSE138746 |
| *MARCO* | 3.5x | 38% | 0.75 | GSE138746 |
| *MERTK* | 3.4x | 45% | 0.60 | GSE138746 |
| *MYOF* | 2.8x | 50% | 0.68 | GSE138746 |
| *NAIP* | 3.3x | 45% | 0.66 | GSE138746 |
| *NFKBIA* | 3.8x | 37% | 0.55 | GSE114407 |
| *NFKBIZ* | 2.7x | 33% | 0.60 | GSE114407 |
| *NLRC4* | 3.0x | 31% | 0.74 | GSE138746 |
| *NR4A1* | 2.5x | 149% | 0.82 | GSE138746 |
| *NRG1* | 4.4x | 55% | 0.85 | GSE138746 |
| *PFKFB3* | 3.2x | 40% | 0.68 | GSE114407 |
| *RHOB* | 3.8x | 66% | 0.72 | GSE138746 |
| *RNF144B* | 2.8x | 48% | 0.75 | GSE138746 |
| *RPH3A* | 4.0x | 66% | 0.73 | GSE138746 |
| *SCO2* | 2.9x | 48% | 0.66 | GSE138746 |
| *SGK1* | 3.4x | 71% | 0.82 | GSE138746 |
| *SHTN1* | 3.6x | 32% | 0.65 | GSE138746 |
| *SIGLEC1* | 3.8x | 171% | 0.94 | GSE138746 |
| *SULT1A1* | 2.5x | 38% | 0.80 | GSE138746 |
| *TCN2* | 4.0x | 51% | 0.65 | GSE138746 |
| *TLR7* | 2.6x | 46% | 0.66 | GSE138746 |
| *TMEM176A* | 3.9x | 73% | 0.92 | GSE138746 |
| *TMEM176B* | 3.9x | 72% | 0.97 | GSE138746 |
| *VNN1* | 4.2x | 63% | 0.80 | GSE138746 |
| *WARS1* | 2.5x | 52% | 0.72 | GSE138746 |

The seven best discriminating monocyte DIRECT LS-TA were then evaluated in the replication samples for group-wise difference and area-under-curve (AUC) in receiver-operator curve (ROC) analysis.

## 3. Results

### 3.1 Shortlisting of potential monocyte cell-type specific (informative) genes

More than 50 monocyte cell-type specific (informative) genes were shortlisted. Monocytes produce >50% of transcripts in PBMC samples as evidenced by achieving the X50 criteria which requires 2.5x folds higher expression in the isolated monocytes than PB samples. These monocyte cell-type specific genes span various cell signalling pathways including genes like *IFI44L*, *IL1B*, *VNN1* and *NFKBIZ*.

The monocyte cell-type specific (informative) reference genes were selected based on a low level of between-sample variance in the monocyte samples by evaluation of coefficient of variation (CV%). That is these reference genes have to fulfilled two criteria at the same time: (1) differential expression criteria for monocyte cell-type specific (informative) genes (X50 criteria), and (2) having the least biological variation among individuals. Among the monocyte informative genes, *PSAP* and *CTSS* has the least CV% at 11% and 9%, respectively. So they were used as the denominator in the monocyte cell-type specific ratio-based biomarker (RBB) or monocyte DIRECT LS-TA in PB.

### 3.2 Correlation between Monocyte Direct LS-TA assays (RBBs in cell-mixture sample) and gene expression of isolated monocytes

The performance of using *PSAP* or *CTSS* as the reference gene (denominator of the RBB) in DIRECT LS-TA was evaluated in those datasets containing gene expression data of both isolated monocytes and cell-mixture samples (PBMC). Gene expression levels in the isolated monocyte samples are the gold standard in this analysis. The ability of DIRECT LS-TA (RBB) assays in PBMC to reflect the gold standard expression levels in isolated monocytes were evaluated by Pearson’s correlation of the 2 expression results. A correlation coefficient (r) > 0.7 was defined as a good correlation between the 2 gene expression results (i.e. R^2^>0.5).

Figure 2 shows a typical workflow of this correlation evaluation between the 2 gene expression results (isolated monocyte and DIRECT LS-TA in PB). For example, *LYZ* is selected as the target gene and its expression level in monocytes is of interest. The gold standard *LYZ* expression level is that in the isolated monocytes and was normalised to a conventional housekeeping gene (*B2M*). The results were plotted on the x-axis as log (*LYZ* _(monocytes)_ /*B2M*_(monocytes)_), equivalent to log *LYZ* _(monocytes)_ - log *B2M* _(monocytes)_ which represented the ground-truth monocyte specific expression level of *LYZ*. The new DIRECT LS-TA (RBB) was determined in the corresponding WB samples of the same individual to evaluate if monocyte specific expression could be readily determined from WB. The DIRECT LS-TA assay of *LYZ* is a RBB using gene expression data in the WB, i.e. log (*LYZ*_(WB)_ / *CTSS*_(WB)_) or equivalent to log *LYZ*_(WB)_ -log *CTSS*_(WB)_. In this example, *CTSS* was used as the monocyte informative reference gene and as the denominator gene in the new RBB in the WB (cell-mixture) sample.

**Figure 2.**
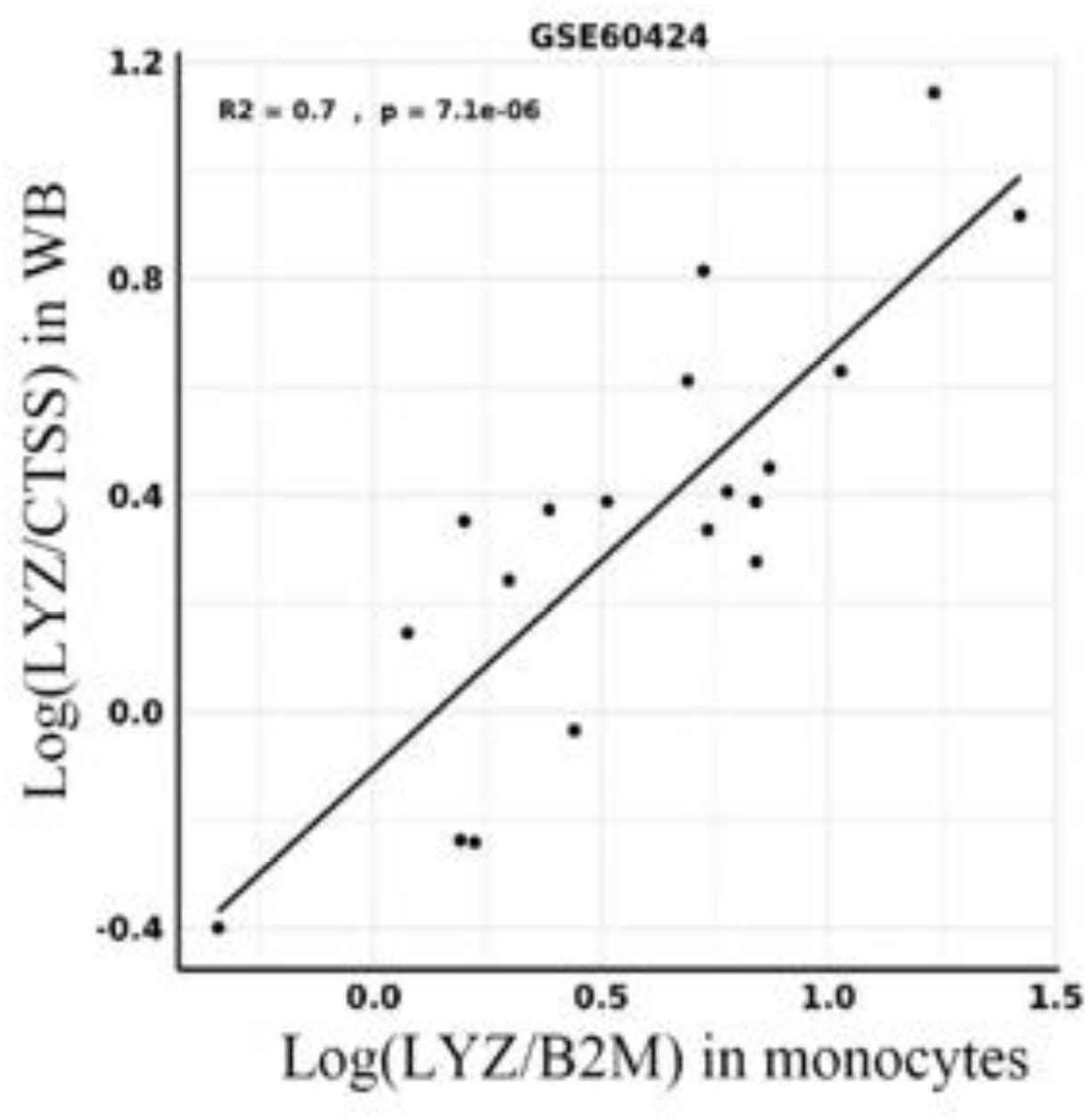
Correlation between the results of biomarkers obtained by the monocyte DIRECT LS-TA assay and the expression of target gene *LYZ* detected in isolated monocytes by the traditional method in the GSE60424 dataset. *LYZ* which is highly expressed genes in monocytes, are selected as the target genes, *CTSS* identified in the present study is used as reference gene in the cell-mixture sample. The X-axis shows the determination of gene expression of *LYZ* in isolated and purified monocytes, using *B2M* as a conventional housekeeping gene, i.e. Log (*LYZ* _(monocytes)_ /*B2M* _(monocytes)_), where the X-axis here is the gold standard. The Y-axis monocyte DIRECT LS-TA in WB, wherein *CTSS* is used as reference gene, i.e. Log (*LYZ*_(WB)_ / *CTSS*_(WB)_). The correlation was high between monocyte DIRECT LS-TA of *LYZ* in WB with monocyte *LYZ* expression (R^2^=0.7).

As shown in table 3, shortlisted monocyte cell-type specific (informative) genes had good correlation between the 2 gene expression results in at least one dataset. R^2^ (coefficient of determination) ranged from 0.55 to up to 0.97 which indicated the ability of DIRECT LS-TA in PB to reflect those specific gene expression in purified monocytes. The results supported that for these shortlisted target genes, their monocyte specific gene expression can be directly determined in PB without the need of isolation of monocytes. And this RBB could be readily used for the differentiation of disease groups.

For comparison, genes used as typical cell markers in flow cytometry such as *CD14* may not work as well as the monocyte specific reference gene in RBB in PB. For example, the correlation between log (*VNN1*_(PBMC)_ / *CD14*_(PBMC)_) and gold standard *VNN1* expression in monocyte was poor with R^2^ at only 0.29 (Supplment Figure 1A). On the other hand, when applying our DIRECT LS-TA method identified denominator gene *PSAP*, the correlation of log (*VNN1*_(PBMC)_ / *PSAP* _(PBMC)_) with gold standard expression was much higher at 0.77 (Supplment Figure 1B).

Figures 3 and 4 showed the extent of correlation between gold standard monocyte gene expression (X axis) and the DIRECT LS-TA method using gene expression measured in PB (Y axis) using *PSAP* and *CTSS* as denominator genes, respectively. For most genes, results in the GSE138746 were shown. Some target genes were not presented in that dataset or had much higher correlation in another dataset and it was used shown instead.

**Supplement Figure 1.**
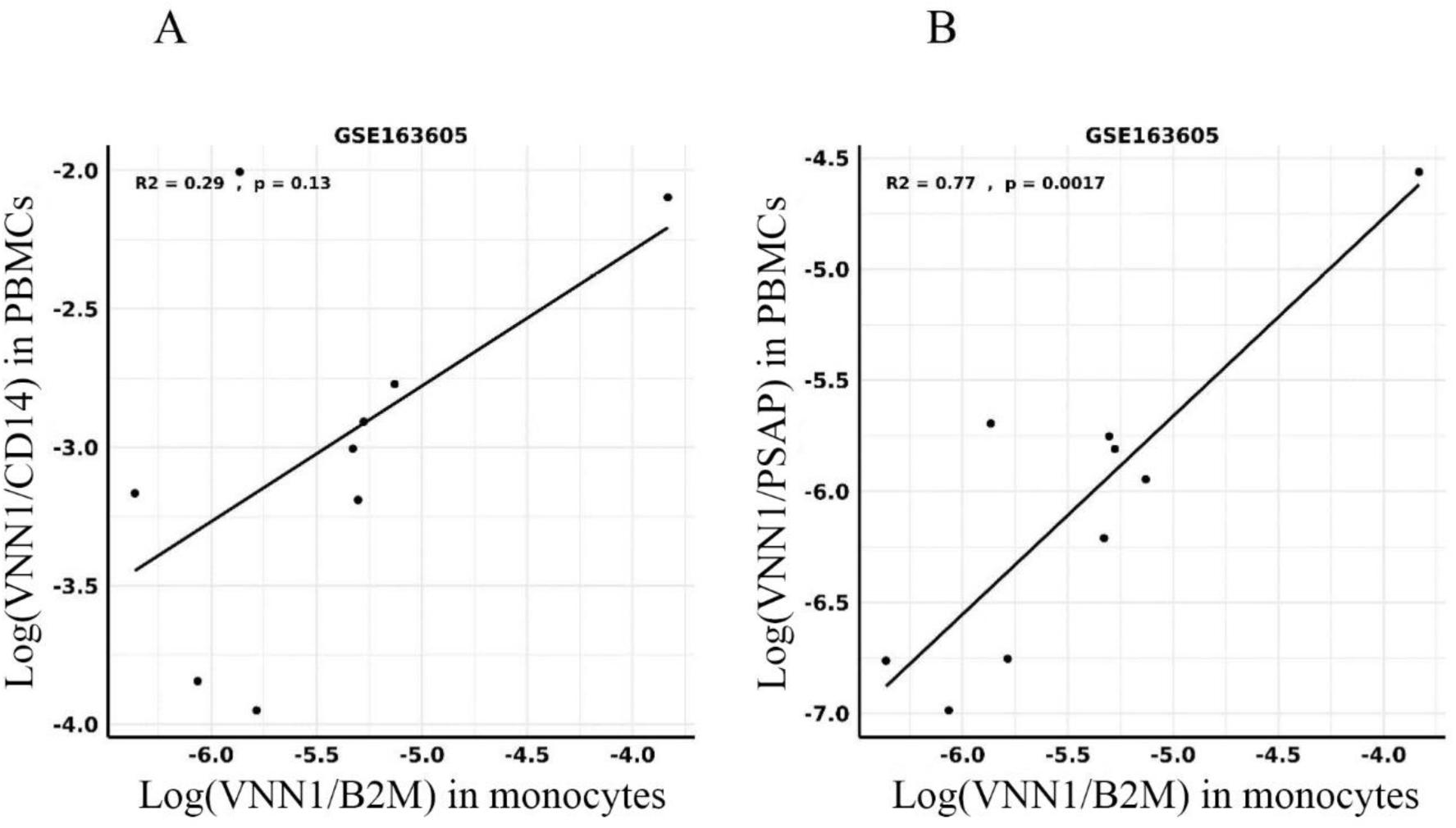
Correlation between the results of biomarkers obtained by the DIRECT LS-TA assay and the gold standard in the GSE163605 dataset. *VNN1*, which is a highly expressed gene in monocytes, is selected as the target gene, and *CD14* and *PSAP* are respectively used as reference genes in the cell-mixture sample. The X-axis shows Log (*VNN1*_(monocytes)_ /*B2M*_(monocytes)_), which is used as the gold standard. The Y-axis shows the RBB in PBMC, wherein *CD14* i.e. Log (*VNN1*_(PBMC)_ / *CD14* _(PBMC)_) and *PSAP* i.e. Log (*VNN1*_(PBMC)_ / *PSAP* _(PBMC)_) are used as reference genes, respectively.

**Figure 3.**
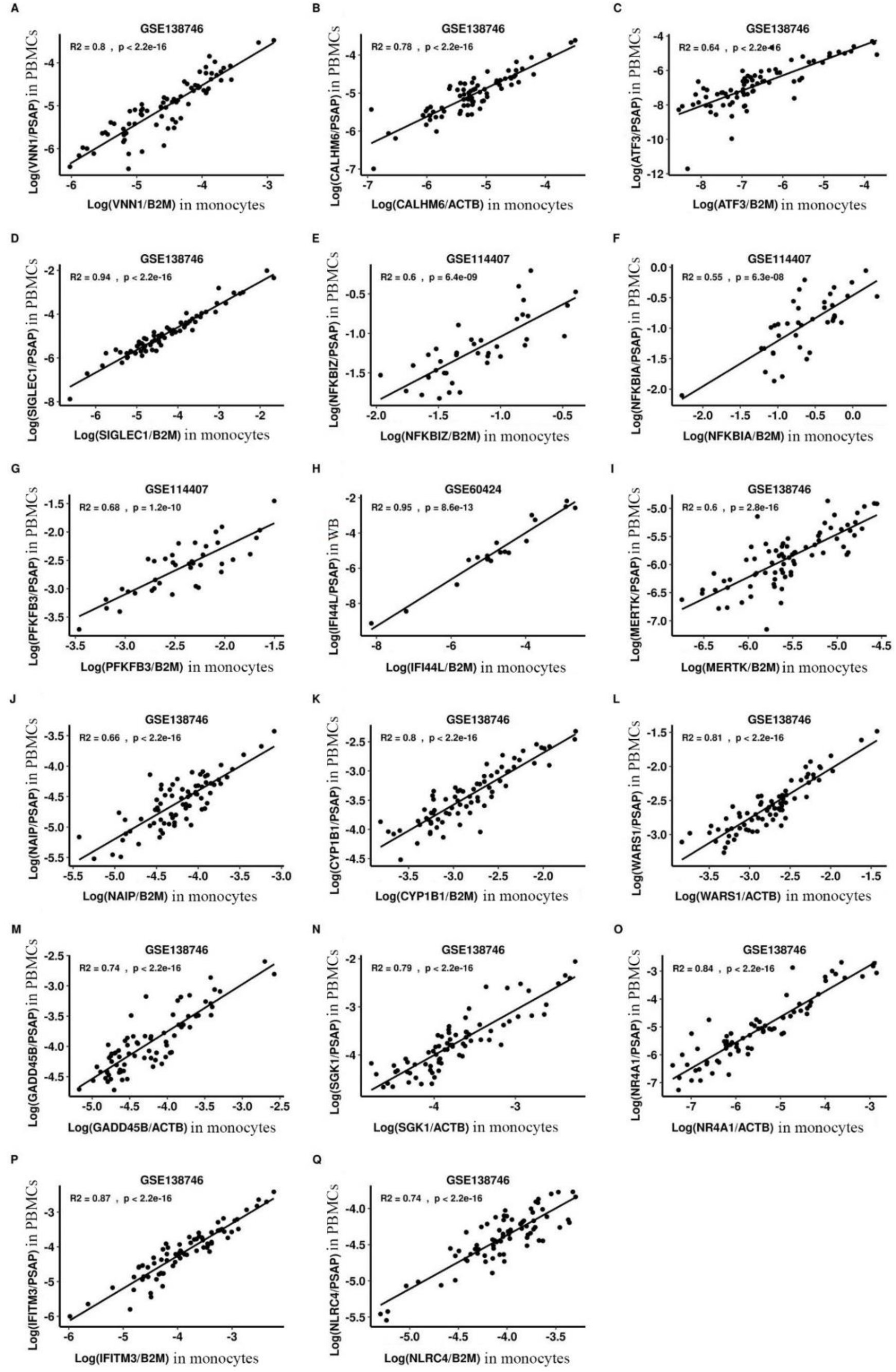
The correlation between monocyte DIRECT LS-TA for monocyte informative target genes measured in the cell-mixture sample of peripheral blood and expression levels of the same target genes in monocytes obtained by the traditional method, using *PSAP* as a monocyte informative reference gene. Figure 3A shows the *VNN1* gene expression in monocytes determined by the method of DIRECT LS-TA and the traditional method. The Y-axis is the ratio of Log (*VNN1*_(PBMC)_ / *PSAP*_(PBMC)_) determined directly from the cell-mixture sample of peripheral blood (i.e., monocyte DIRECT LS-TA biomarker of *VNN1* gene). The X-axis is the gold standard, using the traditional method to detect *VNN1* expression after isolation and purification of monocytes, and using a conventional housekeeping gene (*B2M*) for normalization, i.e. Log (*VNN1*_(monocytes)_ /*B2M*_(monocytes)_). As shown in Figure 3A, there is a good correlation between the two. Evaluation of the performance of other monocyte informative genes using DIRECT LS-TA in peripheral blood is shown in Figure 3B-3Q, where the genes are *CALHM6, ATF3, SIGLEC1, NFKBIZ, NFKBIA, PFKFB3, IFI44L, MERTK, NAIP, CYP1B1, WARS1, GADD45B, SGK1, NR4A1, IFITM3,* and *NLRC4*, respectively. Dataset accession numbers for data sources are shown above Figures 3A-3Q. Logarithms used in Figures 3A-3Q are natural logarithms.

**Figure 4.**
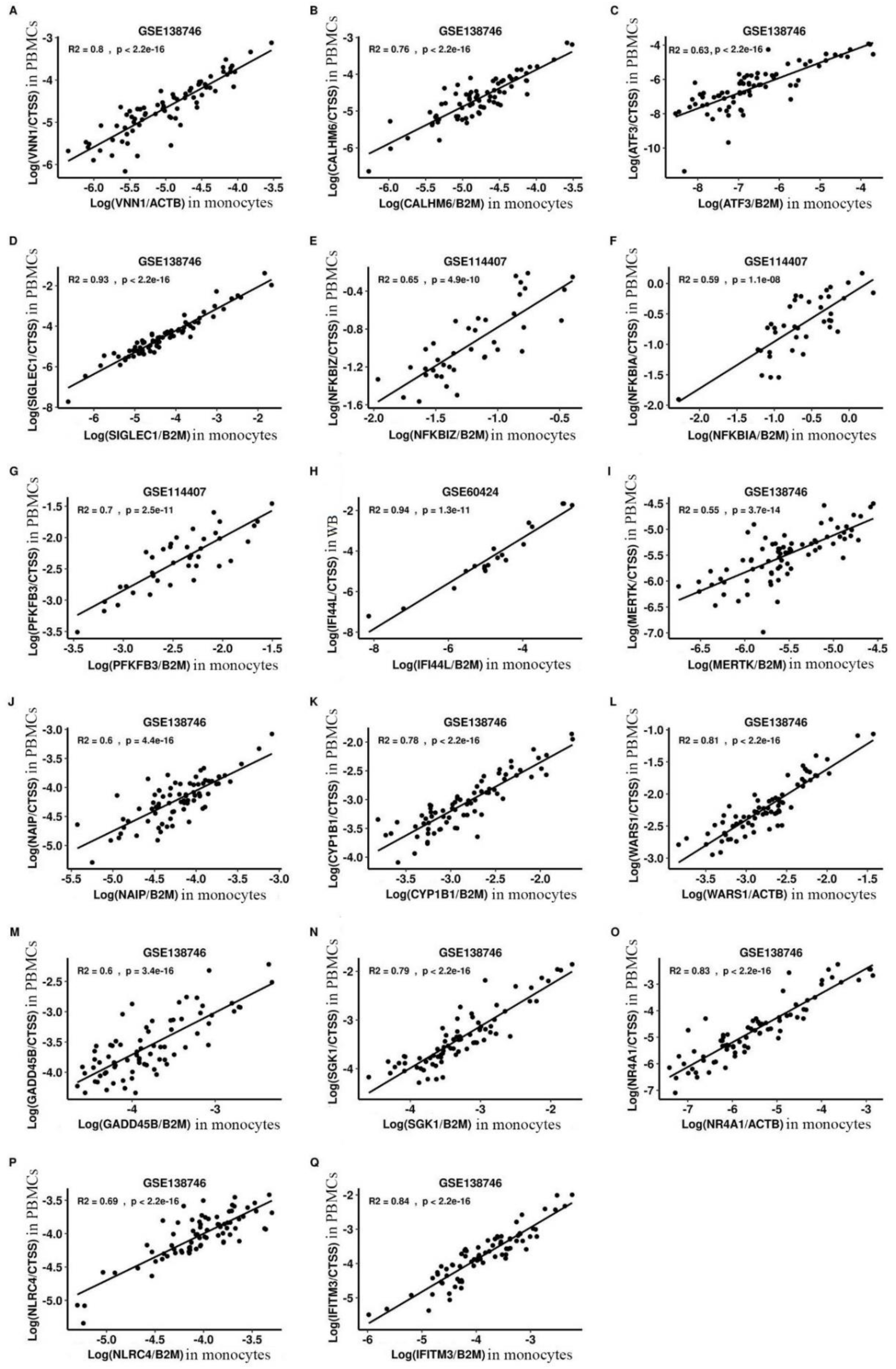
The correlation between monocyte DIRECT LS-TA for monocyte informative target genes measured in the cell-mixture sample of peripheral blood and expression levels of the same target genes in monocytes obtained by the traditional method, using *CTSS* as a monocyte informative reference gene. Figure 4A shows the *VNN1* gene expression in monocytes determined by the method of DIRECT LS-TA and the traditional method. The Y-axis is the ratio of Log (*VNN1*_(PBMC)_ / *CTSS*_(PBMC)_) determined directly from the cell-mixture sample of peripheral blood (i.e., monocyte DIRECT LS-TA biomarker of *VNN1* gene). The X-axis is the gold standard, using the traditional method to detect *VNN1* expression after isolation and purification of monocytes, and using a conventional housekeeping gene (*B2M*) for normalization, i.e. Log (*VNN1*_(monocytes)_ /*B2M*_(monocytes)_). As shown in Figure 4A, there is a good correlation between the two. Evaluation of the performance of other monocyte informative genes using DIRECT LS-TA in peripheral blood is shown in Figure 4B-4Q, where the genes are *CALHM6*, *ATF3*, *SIGLEC1, NFKBIZ, NFKBIA, PFKFB3, IFI44L, MERTK, NAIP, CYP1B1, WARS1, GADD45B, SGK1, NR4A1, IFITM3,* and *NLRC4*, respectively. Dataset accession numbers for data sources are shown above Figures 4A-4Q. Logarithms used in Figures 4A-4Q are natural logarithms.

### 3.4 Direct LS-TA methods to gather monocyte specific gene expression directly in PB and used as biomarkers for bacterial infection

Using the WB gene expression results in GSE154918 dataset (Herwanto et al. 2021), monocyte DIRECT LS-TA of *VNN1* gene is calculated as (Monocyte DIRECT LS-TA of *VNN1*) = (*VNN1*_(WB)_)/(*PSAP*_(WB)_) or its log transformation, log(*VNN1*_(WB)_) - log(*PSAP*_(WB)_). This is similar to delta CT (ΔCT) in qPCR experiments.

To convert to fold change against healthy reference individual, or delta-delta CT (ΔΔCT) in qPCR experiments, MoM of monocyte DIRECT LS-TA is used by subtracting monocyte DIRECT LS-TA results of patients with that of the median in the control group. By setting the median of log Monocyte DIRECT LS-TA of the control (healthy) group to zero, a MoM of log DIRECT LS-TA of each samples is similar to the delta-delta CT values in qPCR or dPCR experiments. It represents the activation (fold change) of target gene over healthy controls in log scale. This is used as biomarker for a disease risk and evaluated for diagnostic performance.

Figure 5 shows the conversion of Monocyte DIRECT LS-TA of *VNN1* from delta CT equivalent (Figure 5A) to delta-delta CT (ΔΔCT) equivalent values (Figure 5B). The sample distribution shown in Figure 5B had no actual change when compared to the sample distribution in Figure 5A. The advantage of using the multiple of median (MoM, Figure 5B) was that the median of the normal control group was set to zero, which allowed comparison of fold changes in DIRECT LS-TA (monocyte specific gene expression) in disease across databases). MoM results can be converted to the expected ΔΔCT results when DIRECT LS-TA is adapted into qPCR or dPCR platform. MoM was 1.2 in Figure 5B which represented a fold change of e^1.2=3.3 fold. The corresponding ΔΔCT results in qPCR is 1.7 cycles.

**Figure 5.**
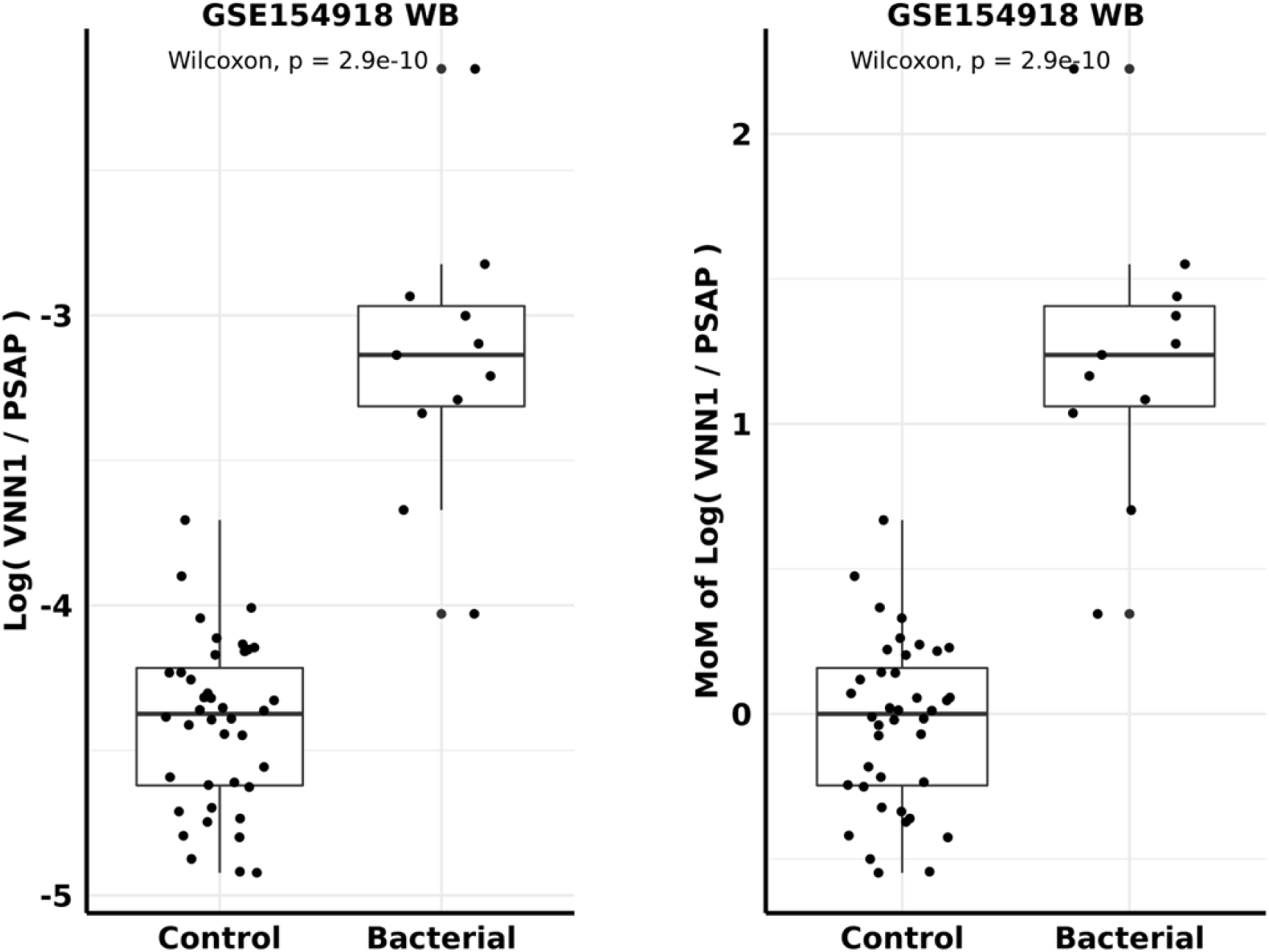
MoM conversion of Direct Monocyte LS-TA of the *VNN1* gene in the GSE154918 dataset. Figure 5A shows the Log (monocyte DIRECT LS-TA of the *VNN1* gene) (i.e., Log (*VNN1*_(WB)_ / *PSAP*_(WB)_) of peripheral blood samples in the control group and the uncomplicated bacterial infection group, respectively. Figure 5B shows the results after converting the Log (*VNN1*_(WB)_ / *PSAP*_(WB)_) in Figure 5A to a multiple of median (MoM). MoM results which is 1.2 in the bacterial infection group. It corresponds to a fold change of 3.3 (e^1.2) and ΔΔCT result of 1.7 cycles when DIRECT LS-TA is adapted into a qPCR platform.

Figure 6 shows the MoM of log (*VNN1*_(WB)_ / *PSAP*_(WB)_) or Monocyte DIRECT LS-TA *VNN1* in healthy controls and bacterial infection patients in the discovery dataset GSE154918 and other 4 replication datasets. Conceptually, the MoM is related to delta-delta CT (ΔΔCT) in qPCR and indicated that there were more than 2 folds increase in Monocyte DIRECT LS-TA *VNN1* in most datasets. GSE60244 had the least activation and the MoM of Monocyte DIRECT LS-TA of bacterial infection patient was ∼1 (natural log), which represented a median increase by 2.72 (e^1) folds. The difference in Monocyte DIRECT LS-TA *VNN1* were highly significant with Wilcoxon non-parametric tests (p values from 2.5×10^−8^ to 1×10^−13^). ROC analysis was also performed on the discovery dataset (AUC=0.99) and in the replication datasets, which returned AUCs ranged from 0.84 to 0.98 (Figure 7).

**Figure 6.**
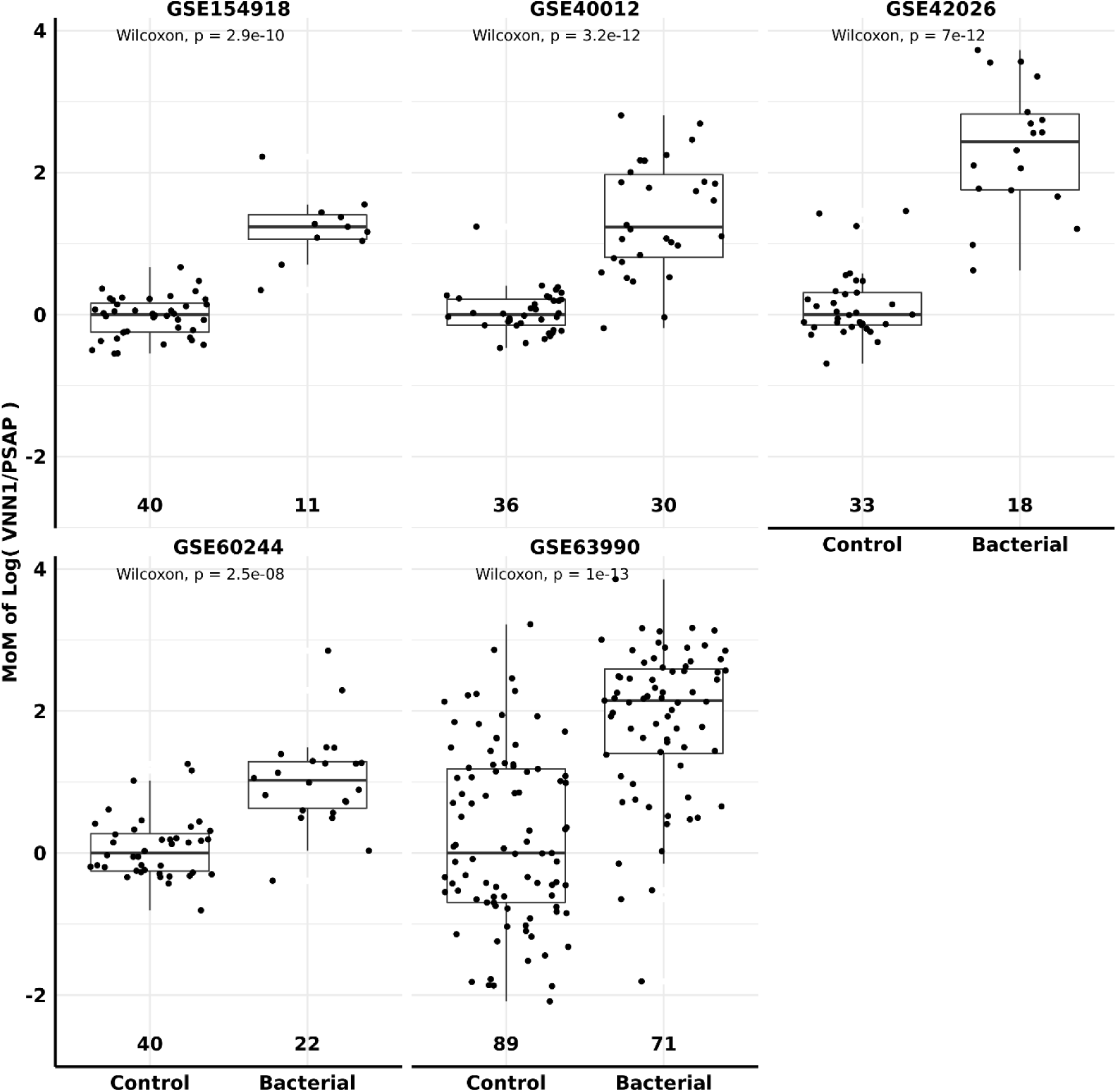
Aanalysis of the MoM of the Monocyte DIRECT LS-TA *VNN1* in the control group and the bacterial infection group. In the five datasets analyzed (GSE154918, GSE40012, GSE42026, GSE60244, and GSE63990, respectively), the numbers above the X-axis represent the number of people in the control group and the bacterial infection group, respectively. In each dataset, the MoM results on the Y-axis are natural log-transformed. MoM results can be converted to the expected ΔΔCT results when DIRECT LS-TA is adapted into qPCR platform.

**Figure 7.**
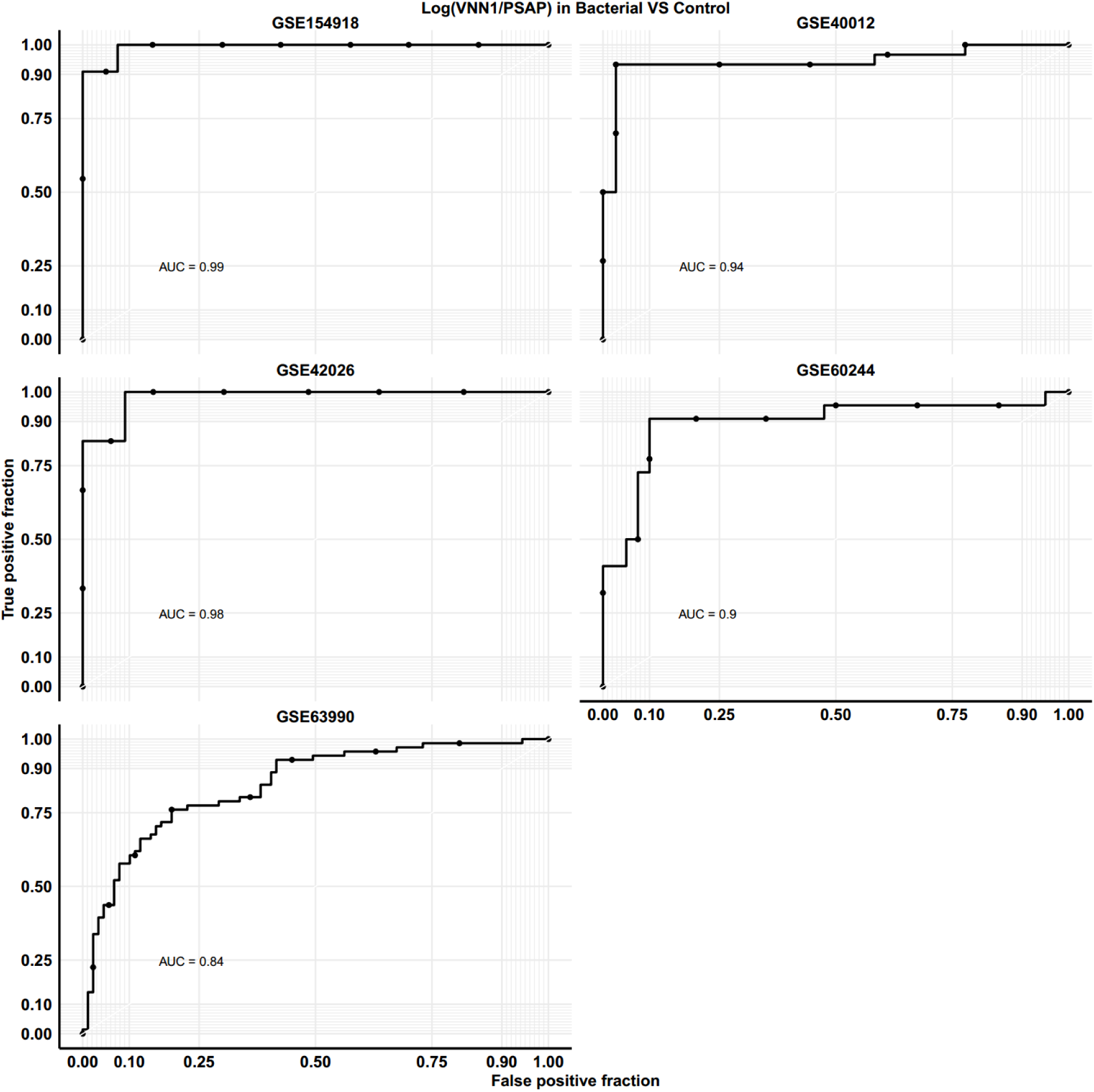
The receiver operating characteristic (ROC) curve analysis of the discriminative performance of the Monocyte DIRECT LS-TA *VNN1* in the bacterial infection group.

Other monocyte informative genes were also activated in patients with bacterial infection including *NLRC4, CYP1B1, PFKFB3, LILRA5, NFKBIA*, and *NFKBIZ*.

Figure 8 showed the MoM of Monocyte DIRECT LS-TA of these additional genes and their diagnostic performance ROC analysis. Wilcoxon group-wise p values ranged from 1.9×10^−6^ (DIRECT LS-TA LILRA5) to 3.8×10^−12^ (DIRECT LS-TA PFKFB3). AUC of MoM of DIRECT LS-TA of these 6 additional target genes were over 0.8.

**Figures 8.**
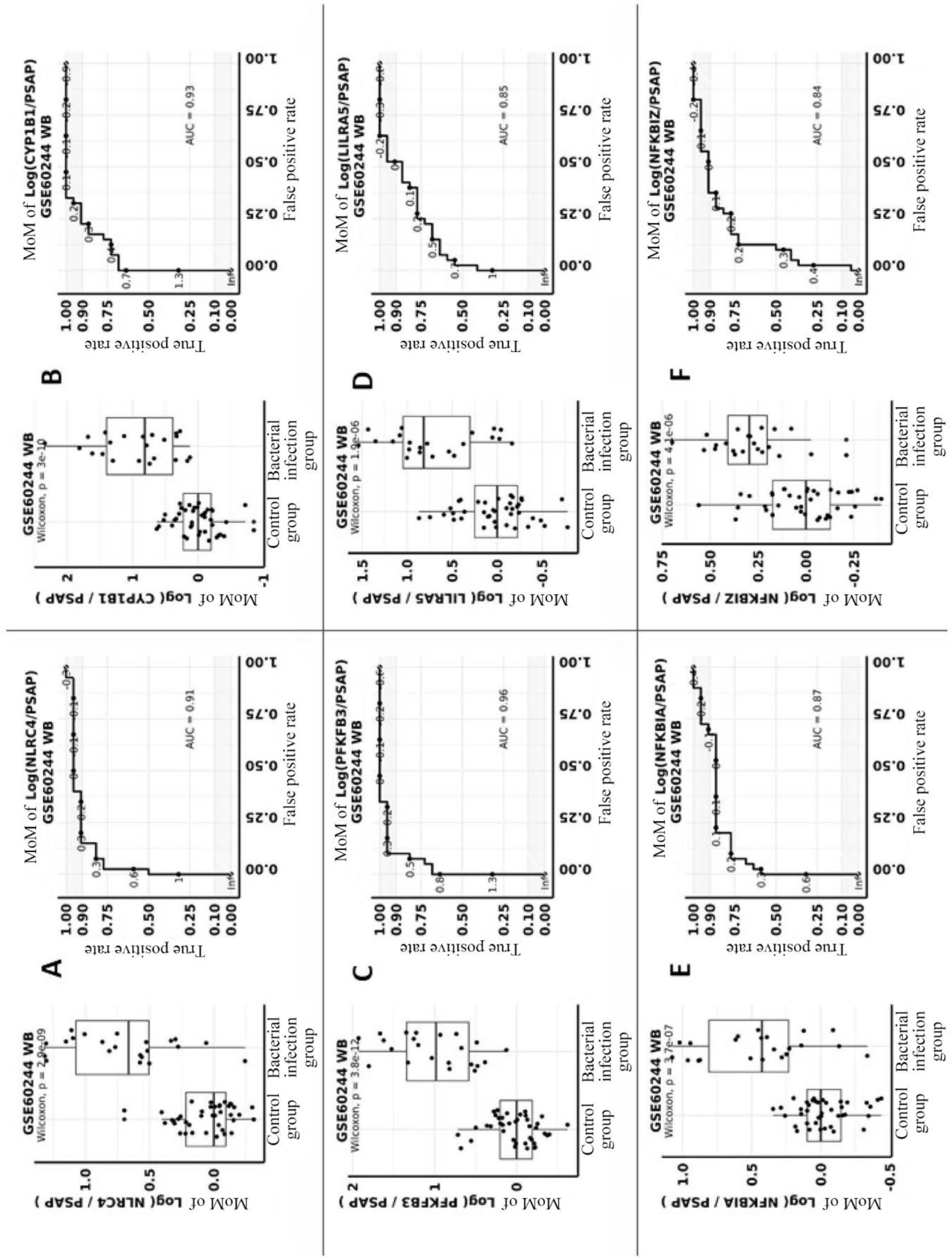
Analysis of the MoM of the Monocyte DIRECT LS-TA for the six additional target gene (i.e., *NLRC4, CYP1B1, PFKFB3, LILRA5, NFKBIA*, and *NFKBIZ*, respectively) using *PSAP* as reference gene in the control group and the bacterial infection group and their receiver operating characteristic (ROC) curve analysis of the discriminative performance in differentiating uncomplicated bacterial infection. Figure 8A-F show the six additional target genes of monocytes, the expression of which are affected by bacterial infection. Here, the Monocyte DIRECT LS-TA is calculated using *PSAP* as a reference gene. For each gene, the difference between MoM of Monocyte DIRECT LS-TA in peripheral blood between the control group and the bacterial infection group is shown by boxplots (left). The right panel shows the ROC for each gene. MoM results can be converted to the expected ΔΔCT results when DIRECT LS-TA is adapted into qPCR platform.

Other than *PSAP*, *CTSS* could be used as the denominator gene of the RBB, Monocyte DIRECT LS-TA. Figure 9 shows the evaluation of Monocyte DIRECT LS-TA using *CTSS* as the denominator gene. Similarly, Wilcoxon group-wise p values ranged from 1.3×10^−6^ (DIRECT LS-TA NFKBIZ) to 6.9×10^−12^ (DIRECT LS-TA PFKFB3). AUC of MoM of DIRECT LS-TA of these 6 additional target genes were over 0.8. The results suggested that both denominator genes (*PSAP* and *CTSS*) produced similar results and performance in term of diagnosis of bacterial infection.

**Figures 9.**
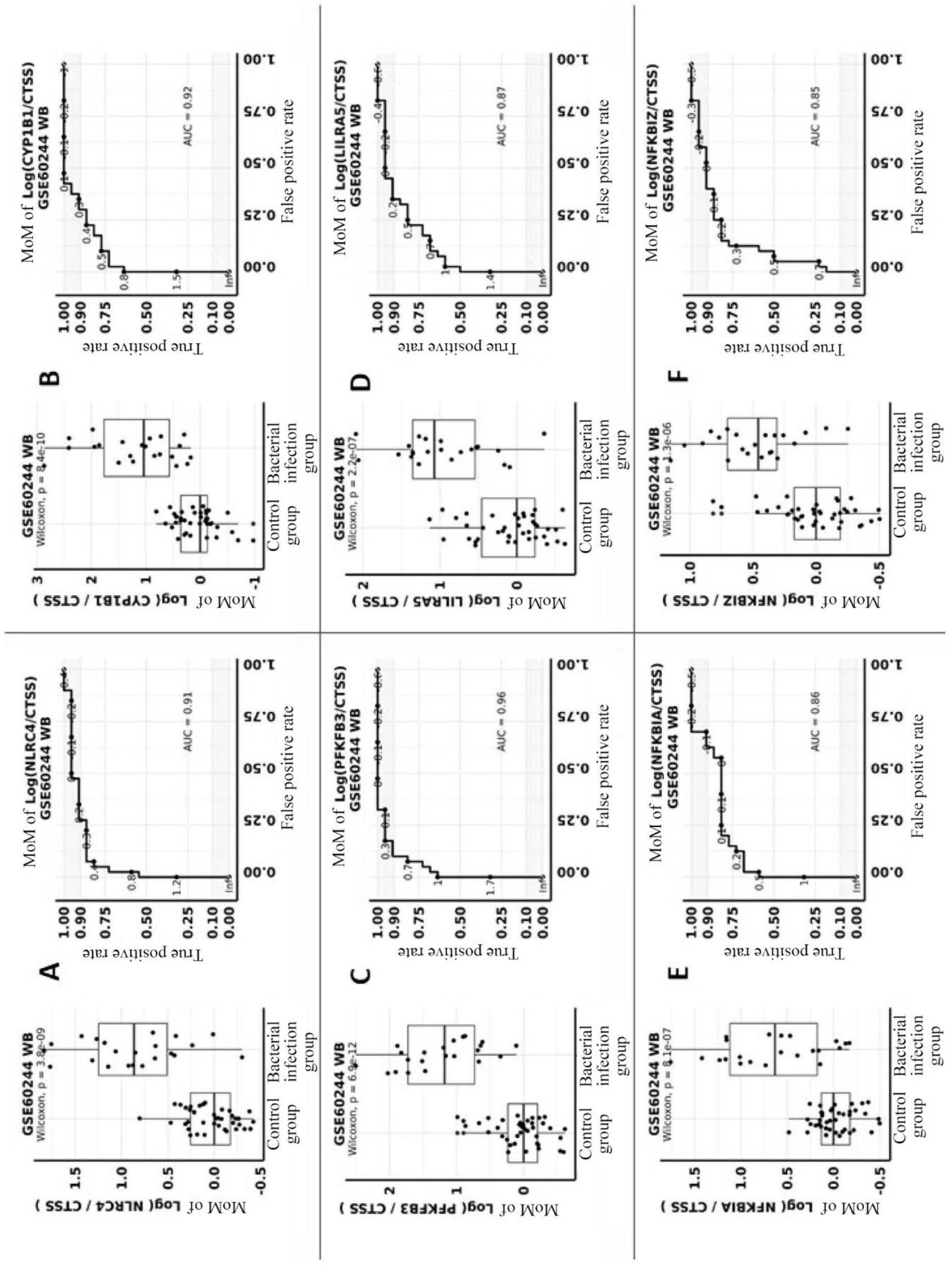
Analysis of the MoM of the Monocyte DIRECT LS-TA for the six additional target gene using *CTSS* as reference gene (i.e., *NLRC4, CYP1B1, PFKFB3, LILRA5, NFKBIA*, and *NFKBIZ*, respectively) in the control group and the bacterial infection group and their receiver operating characteristic (ROC) curve analysis of the discriminative performance in differentiating uncomplicated bacterial infection. Figure 9A-F show the six additional target genes of monocytes, the expression of which are affected by bacterial infection. Here, the Monocyte DIRECT LS-TA is calculated using *CTSS* as a reference gene. For each gene, the difference between MoM of Monocyte DIRECT LS-TA in peripheral blood between the control group and the bacterial infection group is shown by boxplots (left). The right panel shows the ROC for each gene. MoM results can be converted to the expected ΔΔCT results when DIRECT LS-TA is adapted into qPCR platform.

## 4. Discussion

Biomarkers of host response to infection have great clinical applications in triage of patients with fever coming to the clinic or emergency department. Early differentiation of patients with potential bacterial infection is important so that they can be managed promptly and necessary samples are collected for bacteriology investigation in time. Traditional bacterial culture takes days to complete. Even the latest state-of-the-art method using ultra-rapid pathogen ID assay takes at least 12 hours (Kim et al. 2024).

Presently, only C-reactive protein (CRP) and procalcitonin (PCT) are in routine clinical use. Both are serum protein markers so they do not convey any cell-type specific host response information, but just represent an overall systemic host response to infection. Therefore, there are overlapping responses to different types of infection. For example, both viral and bacterial infections lead to elevation of CRP, thus in some patients making such differentiation is difficult by using these protein biomarkers.

PB is a cell-mixture sample of leukocytes of various subpopulations or cell-types (e.g. monocyte, granulocytes and lymphocytes). Cell count proportions of these subpopulations are useful in the differential diagnosis of fever. For example, granulocytes percentage increases in bacterial infection and lymphocyte percentage increases in viral infection. However, the between-individual variations of these cell count proportions are large and it is difficult to get cutoff values to make the differential diagnosis (Table 4).

**Table 4.**
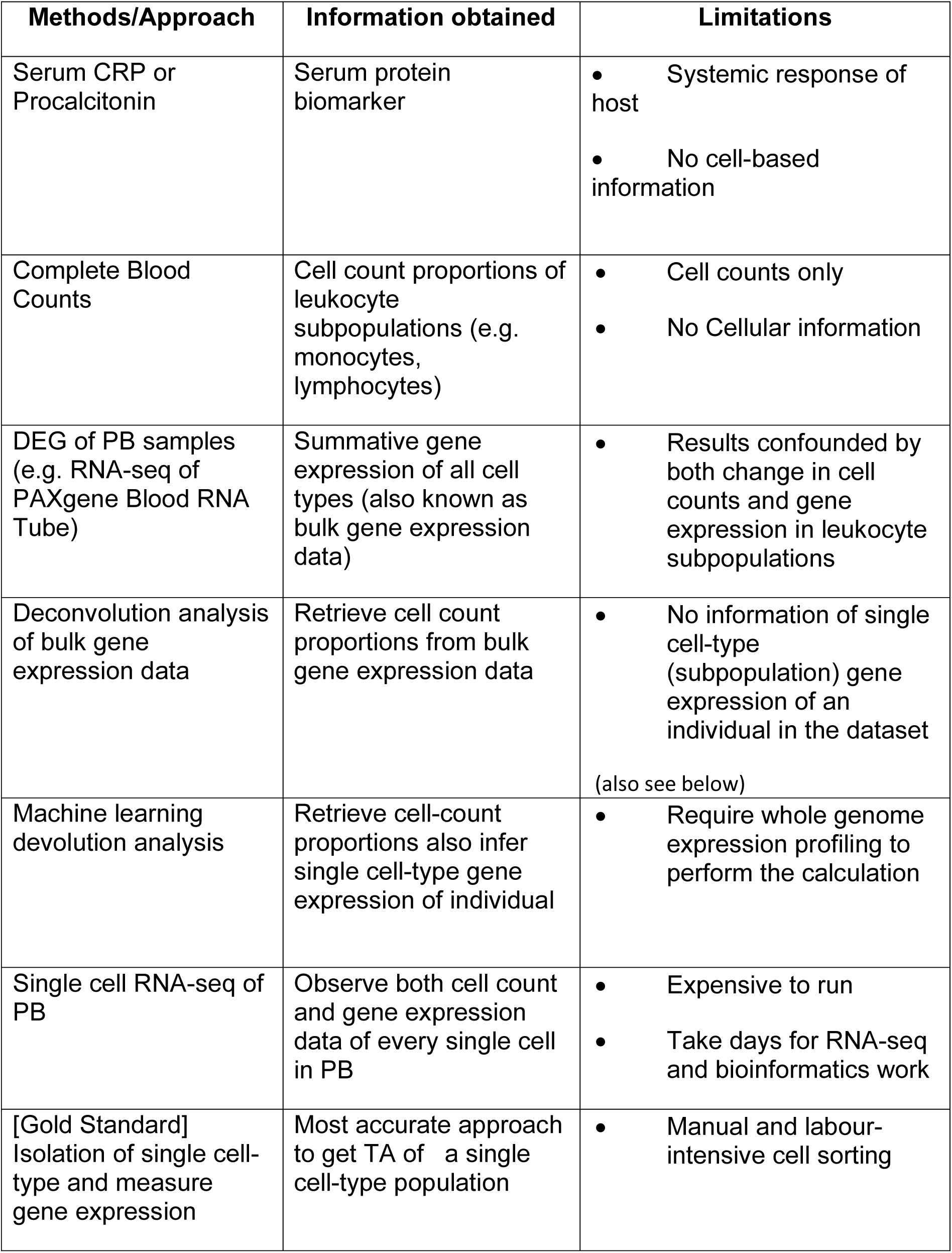
Pros and Cons of Different Approach.

| Methods/Approach | Information obtained | Limitations |
| --- | --- | --- |
| Serum CRP or Procalcitonin | Serum protein biomarker | • Systemic response of host<br>• No cell-based information |
| Complete Blood Counts | Cell count proportions of leukocyte subpopulations (e.g. monocytes, lymphocytes) | • Cell counts only<br>• No Cellular information |
| DEG of PB samples (e.g. RNA-seq of PAXgene Blood RNA Tube) | Summative gene expression of all cell types (also known as bulk gene expression data) | • Results confounded by both change in cell counts and gene expression in leukocyte subpopulations |
| Deconvolution analysis of bulk gene expression data | Retrieve cell count proportions from bulk gene expression data | • No information of single cell-type (subpopulation) gene expression of an individual in the dataset<br>(also see below) |
| Machine learning devolution analysis | Retrieve cell-count proportions also infer single cell-type gene expression of individual | • Require whole genome expression profiling to perform the calculation |
| Single cell RNA-seq of PB | Observe both cell count and gene expression data of every single cell in PB | • Expensive to run<br>• Take days for RNA-seq and bioinformatics work |
| [Gold Standard] Isolation of single cell-type and measure gene expression | Most accurate approach to get TA of a single cell-type population | • Manual and labour-intensive cell sorting |

With the advance in molecular techniques to quantify gene expression, many researchers analyzed the TA in PB by microarray or RNA-sequencing. The expression of each gene is statistically analyzed one by one, and then the genes with the greatest expression difference between different groups were identified as the biomarker. These gene expression biomarkers are also called differential expression genes (DEGs) (Lydon et al. 2019; McClain et al. 2021; Sweeney et al. 2016; Tsalik et al. 2016; Tsao et al. 2020).This method ignores the confounding factor of the cell counts of various cell subpopulations and their variations in different diseases. Therefore, variations in these factors will weaken the effectiveness of DEG biomarkers in differentiating diseases. These studies showed that bacterial infection induced expression of a large battery of genes in PB. Both innate differences in gene expression of various leukocyte cell-types and changes in cell count proportions contribute to confound the identification of DEG biomarkers in diseases. Typically, a long list of genes are needed to differentiate patient groups (Mahajan et al. 2016; Mejias et al. 2013; Zaas et al. 2013). Also, these methods can only be carried out in a research laboratory setting and require expensive equipment(Holcomb et al. 2017). DEG confounded by cell count proportions in the cell-mixture sample of PB is not the optimal IVD to read out the host response.

Computation algorithms have developed to deconvolute the cell-count proportion of each cell type presented in a PB sample using matrix deconvolution (Newman et al. 2015; Shen-Orr et al. 2010). However, these algorithms assumed the same expression profile for each cell-types for all subjects in a group. Only recently, methods are developed to determine single cell-type gene expression for individual subjects in a dataset using machine learning methods (Khatri et al. 2024). However, all these methods require input of the gene expression data of the whole genome such as microarray data or RNA-sequencing data. And these technologies are not ready for everyday clinical use or still expensive to use at the moment. Of course, the gold standard approach is to isolate monocyte from PB and measure gene expression of the target genes. However, it requires cell separation which is the cumbersome and technically challenging in a clinical laboratory setting. It is not practical to implement cell isolation procedures in routine hospital laboratory for the time being. The Pros and Cons of various methods are shown in table 4.

In contrast, by using DIRECT LS-TA method, gene expression of a single cell-type in PB can be directly determined for every sample. In this study, it is used as a biomarker to differentiate bacterial infection using gene expression data available in the public datasets. This new DIRECT LS-TA biomarker is a RBB reflecting the TA of monocyte specific target genes that can be quantified directly in PB samples. The correlation of DIRECT LS-TA results and TA in isolated monocytes was very strong, R^2^ for some target genes were up to 0.9 or even more. DIRECT LS-TA results represent the average gene expression of a single cell-type, and therefore, is not confounded by the cell count proportions in cell-mixture samples. Furthermore, this RBB method can be readily translational to clinical application as it only requires the use of qPCR or dPCR machines which are widely available nowadays in most clinical laboratories.

Antimicrobial resistance remains a global health challenge and over 4 million deaths were estimated to associate with bacterial microbial resistance in 2021(Naghavi et al. 2024). Accurate and timely discriminating diagnosis of bacterial infection is essential to reduce antibiotic misuse and overuse. In this article, we shortlisted activated target genes in monocytes during acute bacterial infection, including *VNN1*, *NLRC4, CYP1B1, PFKFB3, LILRA5, NFKBIA* or *NFKBIZ*. These monocyte informative target genes and *PSAP* or *CTSS* as the monocyte specific reference gene can be used as a new kind of RBB. Such monocyte DIRECT LS-TA assays are useful in differentiating bacterial infection. The high correlation of gene expression of these target genes in isolated monocytes and direct measurement of TA in PB without the need of cell sorting are unique features of Monocyte DIRECT LS-TA method. This technology is feasible to apply in clinical setting to provide a robust and accurate differential diagnosis of bacterial infection.

Our study was limited by confining to use publicly available gene expression datasets and having little control of the design of the original study e.g. case definition of bacterial infection and different platforms of gene expression quantification. Therefore, we confined our case selection to acute uncomplicated bacterial infection and excluded cases with systemic sepsis which is a heterogeneous condition (Herwanto et al. 2021). Also, the sample size of the datasets was small (discovery dataset: 29 bacterial infection patients, Replication datasets: total 87 patients). Moreover, our study and results could only be used to discriminate bacterial infection as a group but not to identify the exact microbial pathogen involved. Other follow-up tests for example blood culture are needed to perform to identify the causative bacteria.

In conclusion, a new and simple peripheral blood biomarker Monocyte DIRECT LS-TA is proposed here which can be readily used in clinical setting. It can be used to differentiate bacterial infection and inform clinicians on the use of antibiotics. DIRECT LS-TA will emerge as a new kind of in vitro diagnostics (IVD) which can convey single cell-type gene expression information from PB samples. The new kind of IVD and uniqueness of the information, together with the ease of implementation will make it very useful in clinics.

## Acknowledgement

The authors declare the following potential conflict of interest. Nelson LS Tang is the inventor of the patent “Determination of gene expression levels of a cell type “which has been assigned to The Chinese University of Hong Kong. K.S. Leung and Nelson LS Tang are share-holders of Cytomics Ltd. Cytomics Ltd. holds a license to use a patent related to DIRECT LS-TA assay. Patent application pending.

